# Xanthine urolithiasis: Inhibitors of xanthine crystallization

**DOI:** 10.1101/335364

**Authors:** Felix Grases, Antonia Costa-Bauza, Joan Roig, Adrian Rodriguez

**Author notes:** Corresponding author: Felix Grases. Laboratory of Renal Lithiasis Research, Faculty of Sciences, University Institute of Health Sciences Research (IUNICS), University of Balearic Islands, Ctra. de Valldemossa, km 7.5, 07122 Palma de Mallorca, Spain.

## Abstract

**OBJECTIVE:** To identify *in vitro* inhibitors of xanthine crystallization that have potential for inhibiting the formation of xanthine crystals in urine and preventing the development of the renal calculi in patients with xanthinuria.

**METHODS:** The formation of xanthine crystals in synthetic urine and the effects of 10 potential crystallization inhibitors were assessed using a kinetic turbidimetric system with a photometer. The maximum concentration tested for each compound was: 20 mg/L for 3-methylxanthine (3-MX); 40 mg/L for 7-methylxanthine (7-MX), 1- methylxanthine (1-MX), theobromine (TB), theophylline, paraxanthine, and caffeine; 45 mg/L for 1-methyluric acid; 80 mg/L for 1,3-dimethyluric acid; and 200 mg/L for hypoxanthine. All crystals were examined by scanning electron microscopy.

**RESULTS:** Only 7-MX, 3-MX, and 1-MX significantly inhibited xanthine crystallization at the tested concentrations. Mixtures of inhibitors had an additive effect rather than a synergistic effect on crystallization.

**CONCLUSION:** Two of the inhibitors identified here —7-MX and 3-MX — are major metabolites of TB. In particular, after TB consumption, 20% is excreted in the urine as TB, 21.5% as 3-MX, and 36 % as 7-MX. Thus, consumption of theobromine could protect patients with xanthinuria from the development of renal xanthine calculi. Clinical trials are necessary to demonstrate these effects *in vivo*.

## Introduction

Xanthine urolithiasis is an infrequent type of renal stone formation caused by xanthinuria, a rare hereditary disorder. An affected patient has a deficiency of xanthine oxidase, resulting in hypouricemia and hypouricosuria (1, 2). Symptom onset can occur at any age. Approximately 50% of patients with classical hereditary xanthinuria present with urinary tract infections, hematuria, renal colic, acute renal failure, and urolithiasis. A small number of patients also develop renal failure, arthropathy, myopathy, or duodenal ulcer (1, 2).

Hereditary xanthinuria is caused by a mutation of xanthine dehydrogenase (XDH, 2p23.1) or molybdenum cofactor sulfurase (MOCOS, 18q12.2) (3-6). These mutations lead to reduced degradation of hypoxanthine and xanthine to uric acid, and the accumulation of these two metabolites. Classically, Type I xanthinuria is caused by an XDH mutation and a deficit of xanthine dehydrogenase/oxidase, whereas Type II xanthinuria is caused by deficits of xanthine dehydrogenase and aldehyde oxidase due to an MOCOS mutation. These different enzyme deficiencies lead to a clinically identical phenotype.

The diagnosis of this disorder is based on plasma and urine levels of uric acid. If an individual has abnormally low values, then clinicians measure urine and plasma xanthine and hypoxanthine levels. Approximately half of these patients have xanthine urolithiasis (7), and some of these patients can progress to renal failure.

The only recommended treatment for patients with xanthinuria is a low purine diet and high intake of fluids. Because the solubility of xanthine is relatively independent of urinary pH, urine alkalinization has no effect (in contrast to patients with uric acid lithiasis) (8,9).

There is a need to identify new agents that can prevent the development of xanthine crystals in the urine of patients with xanthinuria.

## Materials and methods

### Reagents and solutions

Xanthine (X), 1-methylxanthine (1-MX), 3-methylxanthine (3-MX), 7-methylxanthine (7-MX), hypoxanthine (HX), theophylline (TP), paraxanthine (PX), theobromine (TB), caffeine (CF), 1-methyluric acid (1-MU), and 1,3-dimethyluric acid (1,3-DMUA) were purchased from Sigma-Aldrich (St Louis, MO, USA). Synthetic urine compounds were obtained from Panreac (Montcada i Reixac, Barcelona, Spain). Chemicals of analytical/reagent-grade purity were dissolved in ultra-pure deionized water from a Milli-Q system, and passed through 0.45 μm pore filters before use. A xanthine stock solution was prepared daily by dissolving 0.5 g of xanthine in 0.1 L of 1 M NaOH. To avoid precipitation of other compounds, such as calcium oxalate or phosphates, crystallization reactions were performed in a simplified synthetic urine, prepared by dissolving 5.60 g Na_2_HPO_4_·12H_2_O, 2.41 g NaH_2_PO_4_·2H_2_O, and 13.05 g NaCl in 1 L H_2_O.

### Turbidimetric assay

Xanthine crystal formation in synthetic urine and the effects of potential crystallization inhibitors were assessed using a kinetic turbidimetric system. This consisted of a photometer (Metrohm 662), a fiber-optic light-guide measuring cell with an attached reflector (light path: 2 × 10 mm), and a monochromatic light source (550 nm). Crystallization was assessed at constant temperature (37°C) with magnetic stirring (300 rpm).

Synthetic urine (180 mL) was added to a crystallization flask, followed by addition of a xanthine solution (20 mL) to a final xanthine concentration of 500 mg/L. When testing an inhibitor, the desired amount was dissolved in this solution. When the resulting solution reached a temperature of 37° C, then 3.6 mL of 6 M HCl was added to achieve a pH of 6.0 (normal urinary pH), and the timer was switched on. The pH of the final solution was measured at the beginning of each experiment, and the absorbance of the solution (550 nm) was recorded during the entire kinetic assay.

The maximum concentration tested for each of the 10 compounds were: 20 mg/L for 3-MX; 40 mg/L for 7-MX, 1-MX, TB, TP, PX, and CF; 45 mg/L for 1-MU; 80 mg/L for 1,3-DMU; and 200 mg/L for HX.

### Analysis of crystal development

A 100 mL aliquot of synthetic urine containing 400 mg/L xanthine at (pH 6.0) was added to 7 crystallization dishes: 1 without inhibitors, 2 with 20 and 40 mg/L of 3-MX; 2 with 20 and 40 mg/L of 7-MX; 1 with a mixture of 20 mg/L of 3-MX and 20 mg/L of 7-MX; and 1 with a mixture of 20 mg/L of 3-MX and 40 mg/L of 7-MX. The dishes were covered with parafilm, and incubated without shaking at 37°C for 24 h. The crystals were carefully collected, dried, and examined by scanning electron microscopy (SEM).

## Results

We studied the effect of 10 compounds on xanthine crystallization in synthetic urine (Figure 1). Three of these compounds — 1-MX, 7-MX, and 3-MX — significantly inhibited xanthine crystallization, and the others had no effects at the highest tested concentrations (Figure 2). We also examined the possible synergistic effects of these 3 inhibitors. The results indicated additive, not synergistic, effects

**Figure 1:**
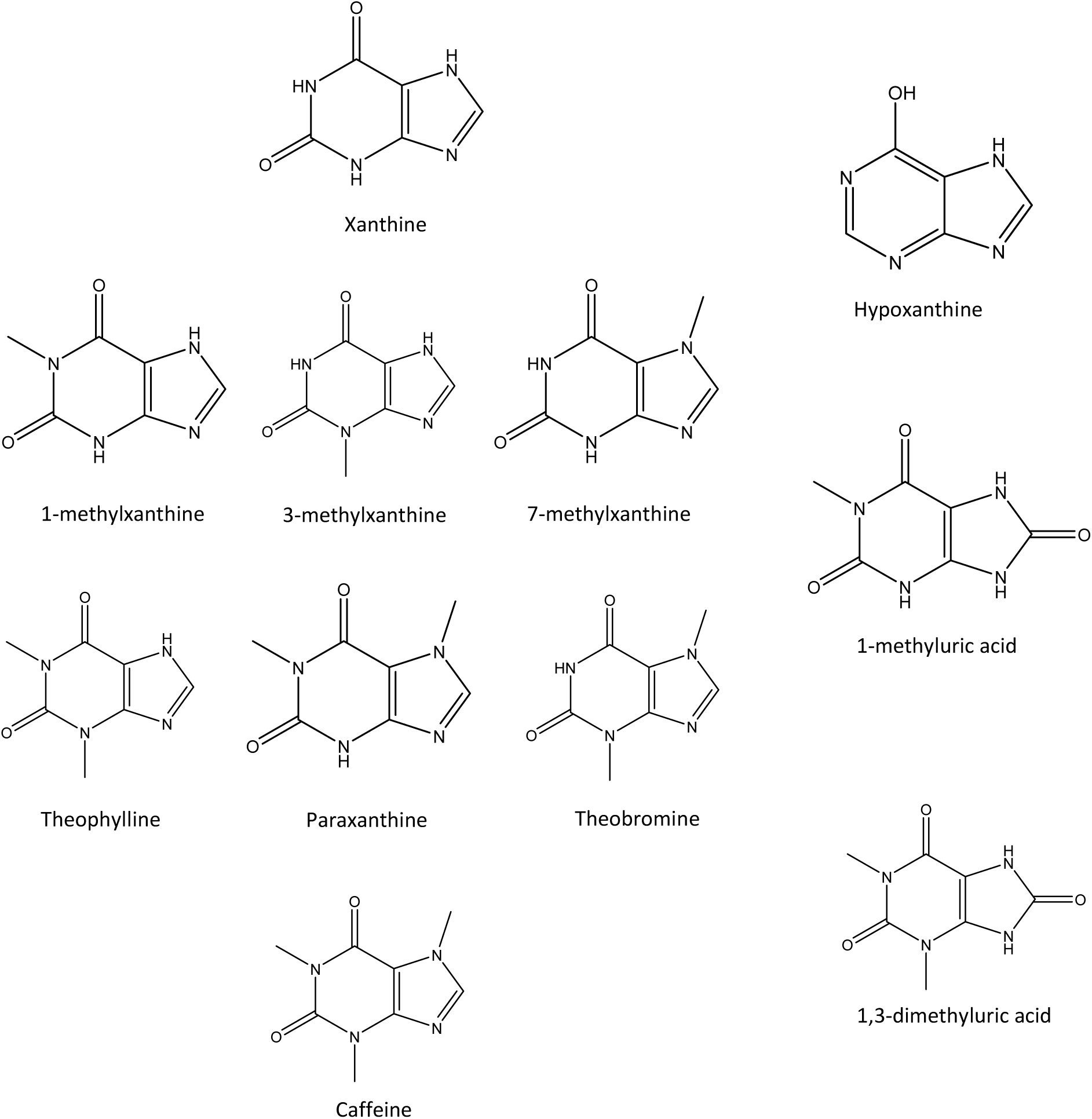
Chemical structure of xanthine and the 10 compounds studied as potential xanthine crystallization inhibitors

**Figure 2:**
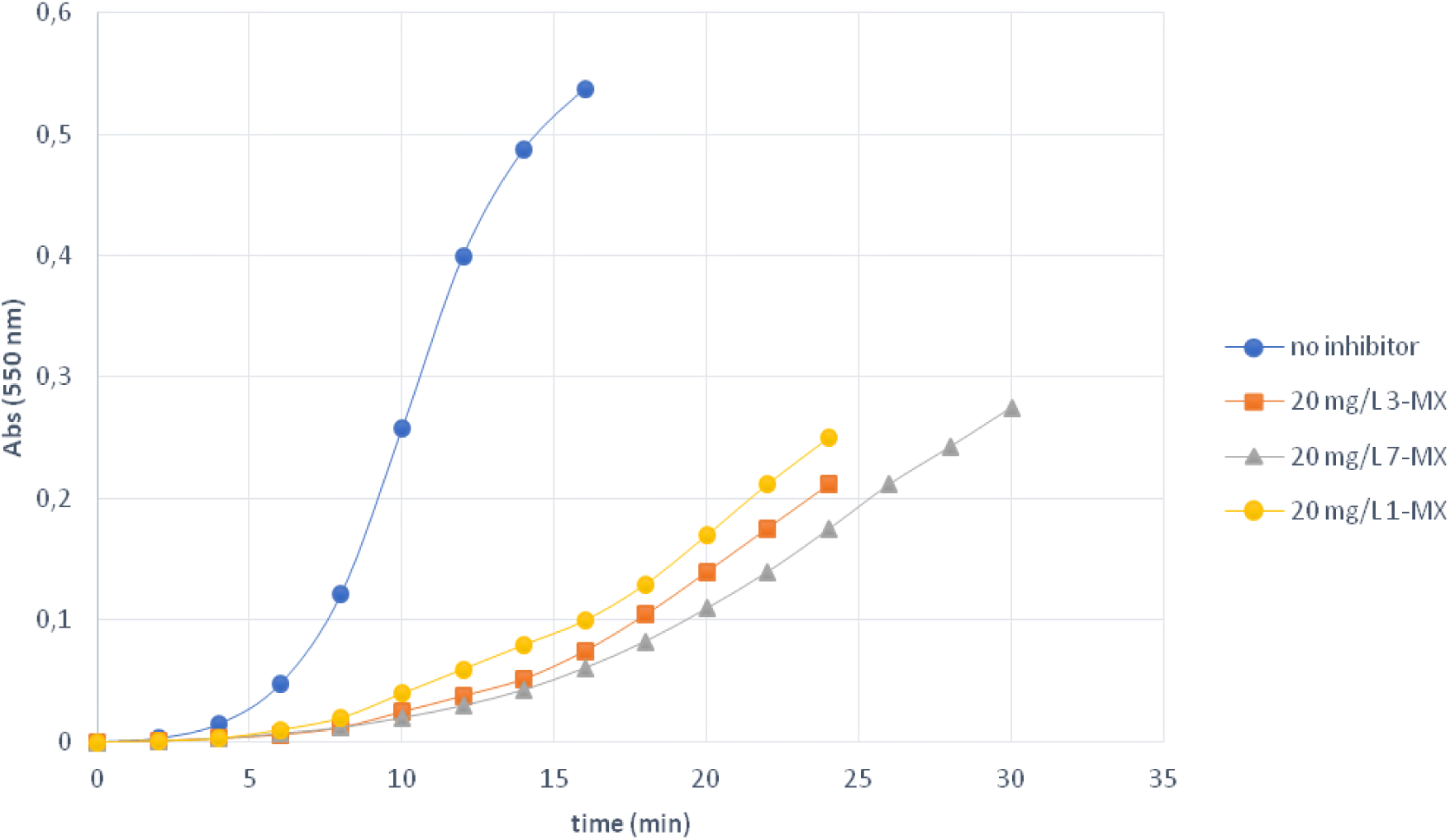
Crystallization curves for 500 mg/L xanthine in synthetic urine, in the absence of inhibitors, and in the presence of 20 mg/L of 1-MX, 3-MX and X mg/L of 7-MX (T=37°C; pH=6.0)

We used SEM to examine the formation of crystals with and without 7-MX, 3-MX, and mixtures of 3-MX and 7-MX (Figure 3). The results show that these inhibitors slowed the rate of crystallization and reduced the abundance of crystalline nucleation; these treatments thereby reduced the formation of crystalline material and large crystals with well-developed faces. Our results clearly demonstrate the efficacy of three inhibitors of xanthine crystallization in artificial urine.

**Figure 3:**
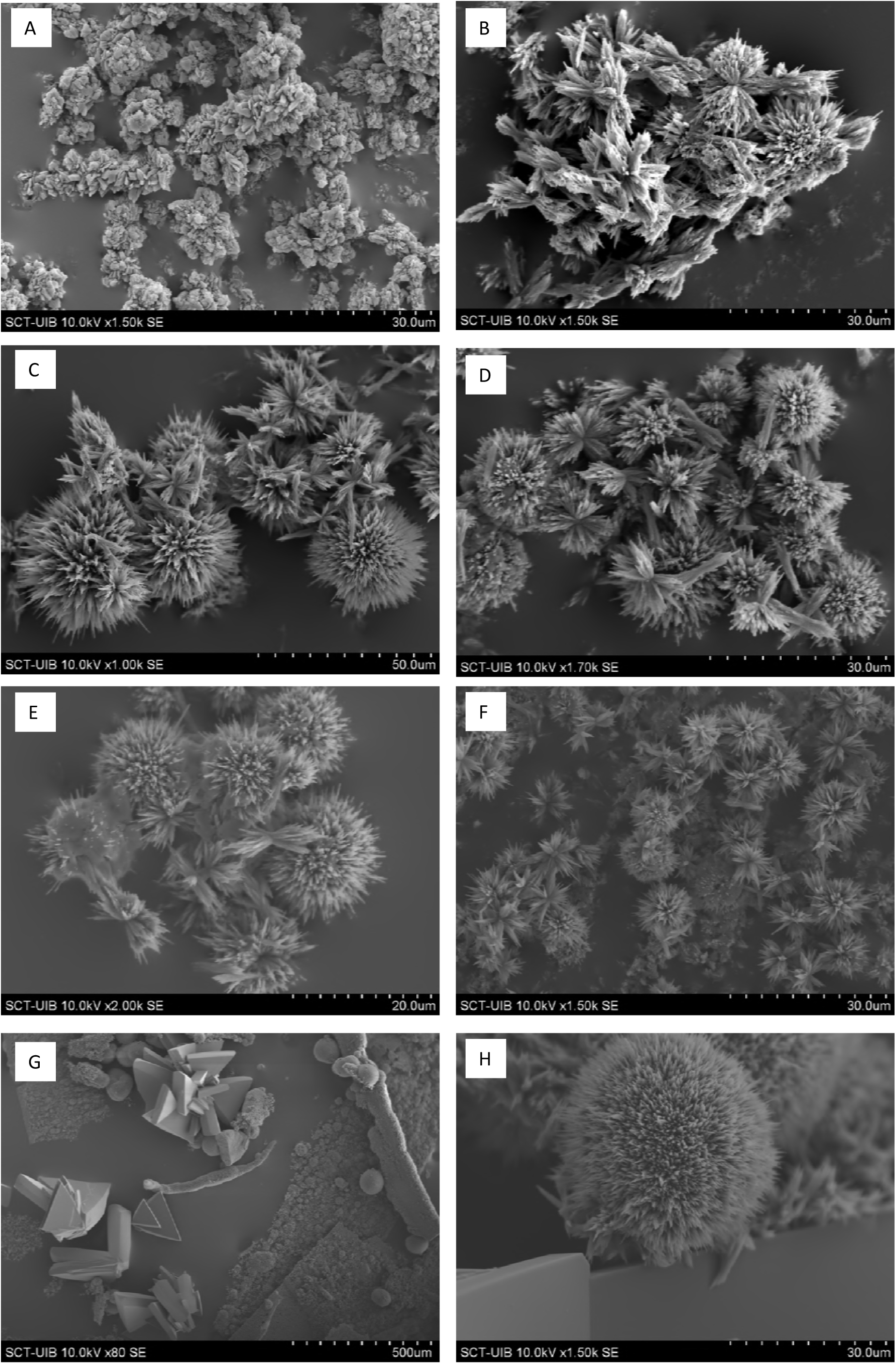
Scanning electron microscopy images of xanthine crystals obtained in synthetic urine containing 400 mg/L xanthine, and incubated 24 h at 37 °C, in the presence of different inhibitors as follows: (A) without inhibitors; (B) with 20 mg/L 3-MX; (C) with 40 mg/L 3-MX; (D) with 20 mg/L 7-MX; (E) with 40 mg/L 7-MX; (F) with 20 mg/L 3-MX and 20 mg/L 7-MX and (G) and (H) with 20 mg/L 3-MX and 40 mg/L 7-MX.

## Discussion

Our results indicate that 1-MX, 7-MX, and 3-MX significantly inhibited xanthine crystallization *in vitro*. Moreover, the delay in time for induction of crystallization was dependent on the concentration of each inhibitor. It is important to note that HX, TP, PX, TB, CF, 1-MU, and 1,3-DMU had no significant effects on xanthine crystallization. This selective inhibition of xanthine crystallization among these structurally similar compounds is notable. Thus, among the xanthines studied, only methyl xanthines are incorporated into the crystalline network of xanthine. The incorporation of any of these 3 inhibitors into the xanthine crystal lattice modifies the structure of some layers, thereby increasing the Gibbs free energy, and slowing the growth of the crystal. Because xanthine solubility is relatively insensitive to urinary pH, we performed all inhibition studies at a pH of 6.0. Interestingly, TB is also very effective inhibitor of uric acid crystallization (10), although CF, PX, and TP have no effect on uric acid crystallization. This demonstrates the importance of the position of the methyl group on the efficacy of an inhibitor.

It is important to consider that two of the inhibitors of *in vitro* xanthine crystallization identified here are major metabolites of theobromine. In particular, after TB consumption, 20% is excreted as TB, 21.5% as 3-MX, and 36% as 7-MX. In contrast, 1- MX is not a metabolite of TB; instead, after consumption of caffeine, 19% is excreted as 1-MX (11). TB is present in high amounts in chocolate and cocoa (12), but has been less studied than other methylxanthines because it has less of an effect on the central nervous system than other xanthines. Extrapolation of previous *in vivo* data (11,12) leads to the estimate that daily intake of only 200 mg of TB would lead to excretion of 43 mg of 3-MX and 72 mg of 7-MX. These amounts, in accordance with the results presented here, could significantly inhibit the formation of xanthine crystals in urine, and therefore prevent the development of renal xanthine calculi in patients with xanthinuria.

Our results indicate that two metabolites of TB — 7-methylxanthine and 3- methylxanthine — can inhibit xanthine crystallization, and therefore have high clinical potential for prevention of nephrolithiasis in patients with xanthinuria. Clinical trials of patients with xanthinuria are necessary to document the *in vivo* effects of TB consumption.

## Acknowledgements

This work was supported by grant PI 14/00853 from the Ministerio de Economía y Competitividad (Gobierno de España), and by FEDER funds (European Union). CIBER Fisiopatología Obesidad y Nutrición (CB06/03), Instituto de Salud Carlos III, Spain, also provided support.

A.R. is grateful to the European Social Fund and the Conselleria d’Educació, Cultura i Universitats for the fellowship FPI/1570/2013.

## References

1.- Carpenter TO, Lebowitz RL, Nelson D, Bauer S, Hereditary xanthinuria presenting in infancy nephrolithiasis. J Pediatr 1986; 109: 309–9.

2.- Mateos FA, Puig JG, Jimenez ML, Fox IH, Hereditary xanthinuria: Evidence for enhanced hypoxanthine salvage. J Clin Invest 1987; 79: 847–52.

3.- Ichida K, Amaya Y, Okamoto K, Nishino T. Mutations associated with functional disorder of xanthine oxidoreductase and hereditary xanthinuria in humans. Int J Mol Sci 2012; 13(11): 15475–95.

4.- Ichida K, Matsumura T, Sakuma R, Hosoya T, Nishino T. Mutation of human molybdenum cofactor sulfurase gene is responsible for classical xanthinuria type II. Biochem Biophys Res Commun. 2001; 282 (5): 1194–200.

5.- Peretz H, Naamati MS, Levartovsky D, Lagziel A, Shani E, Horn I, Shalev H, Landau D. Identification and characterization of the first mutation (Arg776Cys) in the C-terminal domain of the Human Molybdenum Cofactor Sulfurase (HMCS) associated with type II classical xanthinuria. Mol Genet Metab. 2007; 91(1): 23–9.

6.- Yamamoto T, Moriwaki Y, Takahashi S, Tsutsumi Z, Tuneyoshi K, Matsui K, Cheng J, Hada T. Identification of a new point mutation in the human molibdenum cofactor sulferase gene that is responsible for xanthinuria type II. Metabolism. 2003; 52 11): 1501–4.

7.- Raivio KO, Saksela M, Lapatto R. Xanthine oxidoreductase: role in human pathophysiology and in hereditary xanthinuria. In: Scriver CR, Beaudest AL, Sly WS, Valle D (Edds) The methabolic and molecular bases of inheredited diseases, 8th edn. New York. McGraw-Hill., 2001, 2639–52.

8.- Nicoletta JA, Lande MB, Medical evaluation and treatment of urolithiasis. Pediatr Clin North Am 2006; 53: 479–91.

9.- Hisatome I, Tanaka Y, Kotake H, et al. Renal hypouricemia due to enhanced tubular secretion of urate associated with urolithiasis: successful treatment of urolithiasis by alkalinization of urineK+, Na+-citrate. Nephron 1993; 65: 578–82.

10.- Grases F, Rodriguez A, Costa-Bauza A, Theobromine inhibits uric acid crystallization. A potential application in the treatment of uric acid nephrolithiasis; PlosOne 2014, 9 doi: 10.1371/journal.pone.0111184.

11.- Maurice J, Arnaud, Pharmacokinetics and metabolism of natural methylxanthines in animal and man, in: Bertid B. Fredholm (Ed), Methylxanthines, Springer, 2011, New York, pp. 33–91.

12.- Craig WJ, Nguyen TT, Caffeine and theobromine levels in cocoa and carob products 2006; J Food Sci, 49: 302–30.

